# Differential functional connectivity underlying asymmetric reward-related activity in human and non-human primates

**DOI:** 10.1101/2020.01.13.904565

**Authors:** Alizée Lopez-Persem, Léa Roumazeilles, Davide Folloni, Kévin Marche, Elsa F. Fouragnan, Nima Khalighinejad, Matthew F. S. Rushworth, Jérôme Sallet

## Abstract

The orbitofrontal cortex (OFC) is a key brain region involved in complex cognitive functions such as reward processing and decision-making. Neuroimaging studies have shown unilateral OFC response to reward-related variables, however, those studies rarely discussed the lateralization of this effect. Yet, some lesion studies suggest that the left and right OFC contribute differently to cognitive processes. We hypothesized that the OFC asymmetrical response to reward could reflect underlying hemispherical difference in OFC functional connectivity. Using restingstate and reward-related MRI data from humans and from rhesus macaques, we first identified a specific asymmetrical response of the lateral OFC to reward in both species. Crucially, the subregion showing the highest reward-related asymmetry (RRA) overlapped with the region showing the highest functional connectivity asymmetry (FCA). Furthermore, the two types of functional asymmetries were found to be significantly correlated across humans. Altogether, our results suggest a similar pattern of functional specialization between the left and right OFC is present in two primate species.

## Introduction

The orbitofrontal cortex (OFC) is a key brain region involved in complex behavior such as value-based decision-making (1), cognitive flexibility (2) and state space representation (3). This brain region is heterogenous and can be subdivided on the basis of cytoarchitecture, connectivity, or function (4–8). The large majority of studies investigating the functional organization of the OFC consider it to be symmetrically organized between hemispheres (1, 9–12). Some unilateral lesion and stimulation studies have nevertheless shown differential behavioral effects. For instance, direct intracortical stimulation in humans showed a left lateralization of negative experience compared to neutral experience (13). Patients with right OFC lesions were more impaired in the Iowa Gambling Task than those with left lesions (14). Asymmetrical OFC responses in healthy subjects have also been reported in fMRI studies (for meta-analyses, see (15, 16)). However, this result has rarely been discussed.

Lateralization of functions in the prefrontal cortex has been shown previously, in particular for language processing (17), visuo-spatial attention (18), but also for relational integration reasoning (15). In humans, reductions in asymmetry have been associated with impaired cognitive functions (19) and hemispheric specialization is suggested to increase processing abilities by reducing bilateral redundancy (20) indicating that there may be some benefit when homotypical areas in each hemisphere specialize. Lateralization of functions has also been reported in non-human primates in the context of audition and vocalization (21–24), or attention (25). Yet, lateralization in other contexts, such as reward processing, has not received much attention in any species.

Using data from the Human Connectome Project, and data collected in rhesus macaques *(Macaca mulatta*), we assessed the nature of the asymmetrical OFC response during reward tasks. First, we identified an asymmetrical response to reward in a specific area of the OFC in both species. Second, we observed that the connectivity of the OFC with the rest of the brain was significantly different between hemispheres. Interestingly, the brain region responding differentially in the reward task was the same as the brain region showing asymmetrical whole-brain connectivity. Moreover, the two types of functional asymmetry were correlated across individuals. Together, our results suggest that the left and right OFC might support different functions – that remain to be characterized, due to an intrinsic difference in their connectivity to the rest of the brain.

## Results

### Asymmetric reward-related activity in the orbito-medial prefrontal cortex (OMPFC)

#### Humans

We selected 57 subjects from the Human Connectome Project for which rs-fMRI had been obtained at 7T and who participated in a gambling task designed to assess reward processing and decision-making (26). Participants had to guess whether a hidden card was higher or lower than a visible card. They received positive, neutral or negative monetary feedback according to the correctness of the response (see Methods). In the fMRI data, we focused on the contrast ‘Reward versus Punishment’ to localize the reward-related activity in the whole brain (Figure 1A). Replicating previous results from a larger dataset (26), this contrast also revealed higher activity for reward compared to punishment in the ventromedial prefrontal cortex (vmPFC) and in the ventral striatum. Interestingly, a significant cluster was found in the right OFC, but not in the left OFC (cluster-corrected, cluster size > 150 voxels). Note that the uncorrected map did not reveal a response in the left OFC either (Figure 1A).

**Figure 1 –.**
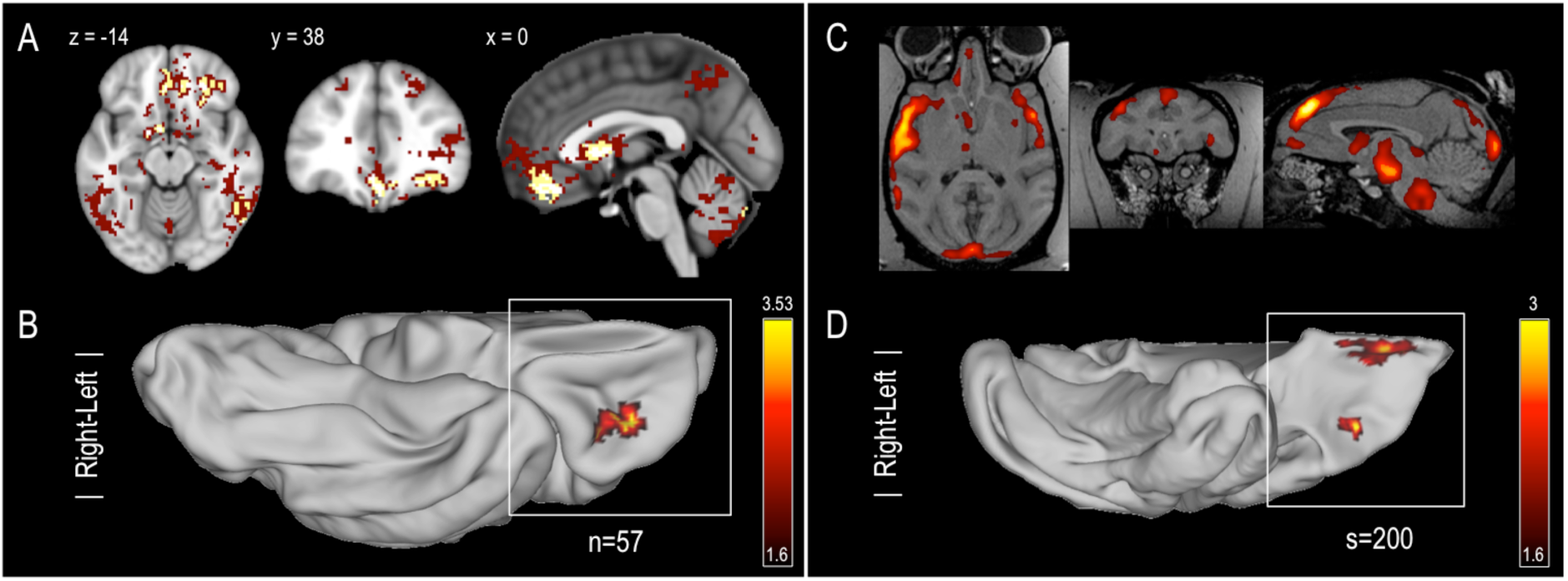
Neural responses to reward and hemispheric differences in reward responses in humans and macaques. **A.** Statistical maps relating to the contrast ‘reward versus punishment’ in humans. Clusters in yellow show significant positive effect (FWE corrected, p<0.05). Clusters in dark red indicate uncorrected effect at p<0.001. **B**. Unsigned difference between the sizes of the effects illustrated in A in the left and in the right hemispheres. Color code indicates z-statistics at the group level, the map is restricted to the OMPFC and cluster corrected (cluster-level p<0.05, permutation tests). **C**. Average of the individual session statistical maps relating to various reward contrasts in macaques (see Methods). Because of the large difference in the number of human and macaque individuals tested, the map is arbitrarily thresholded to illustrate similarity of response with human data. **D**. Unsigned difference between the sizes of the effects illustrated in **C** in the left and in the right hemisphere. Color code indicates z-statistics at the group level, the map is restricted to the OMPFC and show clusters larger than 10 vertices.

To assess whether this hemispherical difference was significant, we mapped the individual z-maps onto the individual MSMAll surfaces, that are registered on the symmetric MNI 152 template (27). We mirrored the data of the left hemisphere so they could be compared to the data on the right hemisphere. We computed the unsigned left versus right difference in the contrast ‘Reward versus Punishment’ for every subject and tested for significant effect at the group level in a large Orbital and Medial Prefrontal Cortex (OMPFC) mask (see Methods). We found a significant difference between left and right OMPFC for reward-related activity in the OFC (p_corr_=0.012) (Figure 1B). This result reveals asymmetric reward-related activity at the intersection of the lateral orbitofrontal sulcus (LOS) and transverse orbitofrontal sulcus (TOS).

#### Macaques

Reward-related asymmetry in macaques was investigated in fMRI data collected from previous studies (see Methods). Eight monkeys who performed different types of reward-related tasks were included in the analyses. For each monkey, we used the reward-related contrasts (see Methods) of each session and averaged them across sessions and individuals to obtain a whole-brain map of reward-related activity (Figure 1C). As in human participants, we projected each session map to a common surface and computed the unsigned left versus right difference in all available contrasts. We found two large clusters (larger than 10 vertices) of reward-related asymmetry (z>2.3) in the OMPFC. First, we observed asymmetric reward-related activity close to the medial orbital sulcus. Second, we also identified a cluster close to the LOS, just posterior to its intersection with the TOS.

In summary, this second area of asymmetry in macaques lies in a similar location with respect to sulcal landmarks in the two species (Figure 1D). In humans it corresponds to the caudal part of area 11l, extending into area a47r according to the parcellation of Glasser et al, 2016 (28). This location corresponds to the caudal part of 47/12m in both humans and macaques in the standard cytoarchitectonic framework proposed by Mackey and Petrides (29).

### Asymmetric functional connectivity in the OMPFC

To determine whether this asymmetry could be explained by an asymmetry in the functional connectivity of the OMPFC, we compared the connectivity profiles of the left and right OMPFC. In both humans (n=57) and macaques (n=14), for each vertex of the OMPFC, we extracted the connectivity (strength of correlation between time series) with all vertices in the brain, from the group-level time series dataset (computed with MIGP, see Methods). The procedure was repeated for the left and the right OMPFC and in each case it was repeated to measure connectivity with ipsilateral and contralateral hemispheres (thereby creating two maps illustrated in figure 2A). The procedure was then repeated a further two times to examine the connectivity of left and right OMPFC with the left hemisphere (regardless of whether the left hemisphere was ipsilateral or contralateral) and the right hemisphere (again, regardless of whether it was ipsilateral or contralateral). It was then possible to assess whether there was any asymmetry in OMPFC connectivity with either the ipsilateral or contralateral hemisphere or with either the left or the right hemisphere. (Figure 2B and C, see Methods). We found in each of the four resulting maps of human OMPFC functional connectivity at least one cluster in the OFC with a particularly high asymmetry. The conjunction of the four maps revealed a unique cluster (Figure 2D). In the following analyses, the FCA measure corresponds to the average of the four types of asymmetry measures. We confirmed the significance of FCA in this cluster at the group level in humans (t(56)=12.29, p=2.10^-17^). The same analysis conducted in macaque data revealed very similar results; the conjunction analysis showed a single cluster in the OFC, with a significant FCA at the group level (t(13)=3.01, p=0.01).

**Figure 2 –.**
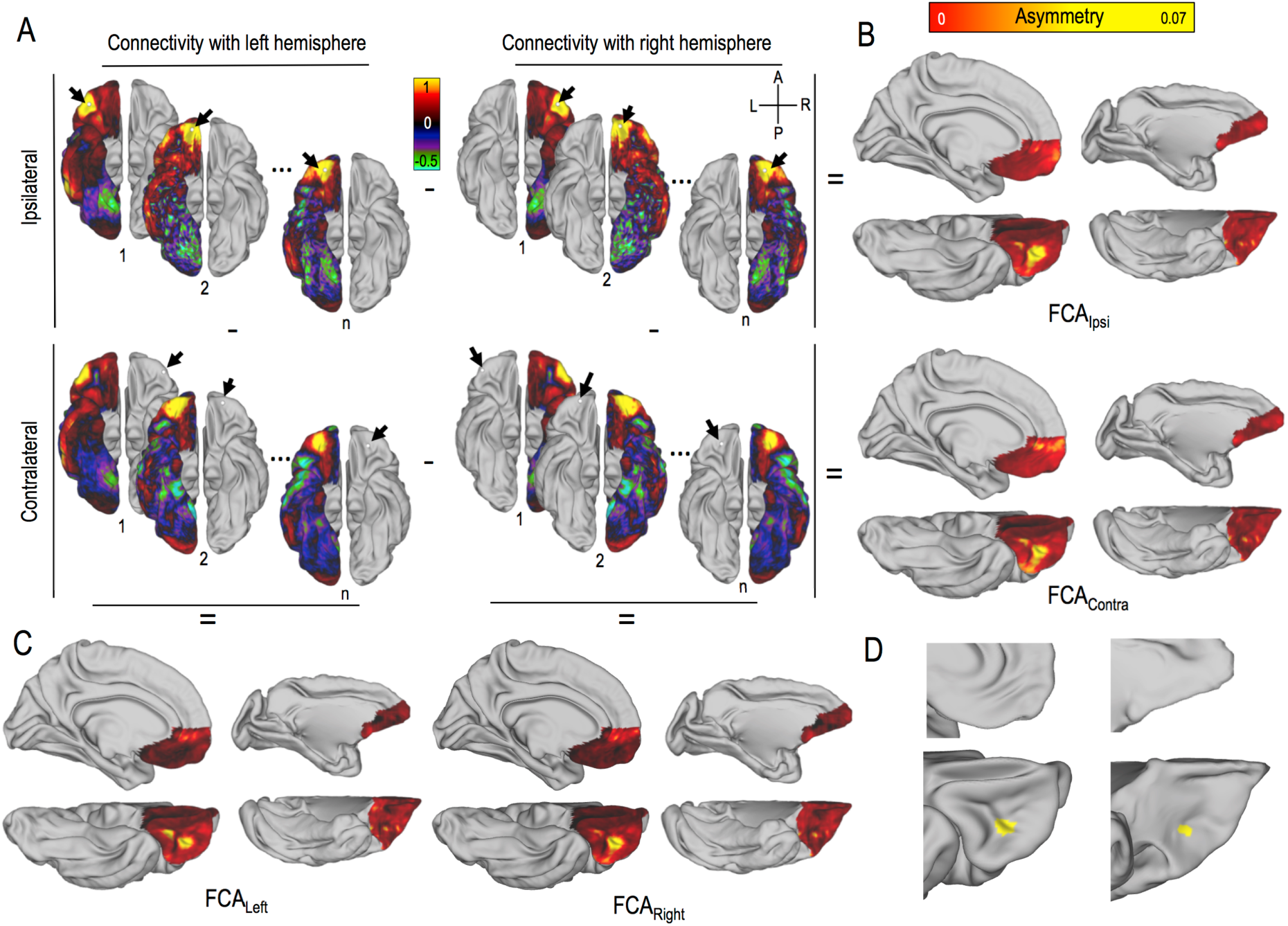
Functional connectivity asymmetry in the human and macaque OMPFC. **A**. Schematic representation of the method to compute FCA measures. **Top row**. Ipsilateral frame. The unsigned difference between the functional connectivity of each vertex in the left OMPFC with all vertices in the left hemisphere and the functional connectivity of each vertex in the right OMPFC with all vertices in the right hemisphere is computed. The l**eft (right) columns** display results for the left (right) hemisphere respectively. Arrows represent the location of seeds while n is the number of vertices in the OMPFC. Colors indicate correlation coefficient between timeseries of the seed and timeseries of each other vertex. **Bottom row**. Contralateral frame. Same as top except that the difference in connectivity is based on the contralateral connectivity of the left and right OMPFC. **B**. The results of these two comparisons between the left and right hemispheres, within the ipsilateral frame (FCA_Intra_) and the contralateral frame (FCA_Contr_) are displayed in the **top row** and **bottom row** for humans (**left**) and macaques (**right**). (**C**) The maps resulting from comparison of left and right OMPFC connectivity with the left (FCA_Left_) and right (FCA_Right_) hemispheres regardless of whether the hemisphere is contralateral or ipsilateral to the OMPFC region examined. Again humans are shown on the **left** and macaques are shown on the **right**.. Hot colors in B and C indicate high asymmetry in functional connectivity. (**D**) Each map in B and C was then z-scored, thresholded (z>2.3), and clusters surviving correction for multiple comparisons were overlapped in humans (left) and macaques (right). Conjunction analyses of the 4 measures of asymmetry revealed the same cluster of functional asymmetry. In panels B, C, and D results are summarized on left surfaces: A, L, R, P corresponds to Anterior, Left, Right, Posterior respectively

### A hotspot of asymmetry in the OFC

#### Overlap between reward-related cluster and functional connectivity cluster

To examine the link between reward-related asymmetry and functional connectivity asymmetry, we projected the results from the two previous sets of analyses onto a common surface (Figure 3). We observed partial overlap of the two clusters in the lateral OFC, in both humans and macaques, indicating unique hotspots of functional asymmetry, as defined by both reward-related activity and by functional connectivity, in the OFC in both species. We computed the coefficient of functional connectivity asymmetry (FCA) in the reward-related asymmetry (RRA) lateral clusters and found that it was significantly higher than in the rest of the OMPFC (humans: t(56)=13.4, p=4.10^-19^, 14 macaques with rs-MRI: t(13)=6.03, p=4.10^-5^, medial cluster: t(13)=-0.51, p=0.62). The reverse analysis, i.e. the investigation of the response difference to reward-related activity in the FCA cluster also revealed a significant response difference to reward in the left and right OFC (humans: t(56)=4.3, p=7.10^-5^, macaque contrasts: t(17)=2.64, p=0.017).

**Figure 3 –.**
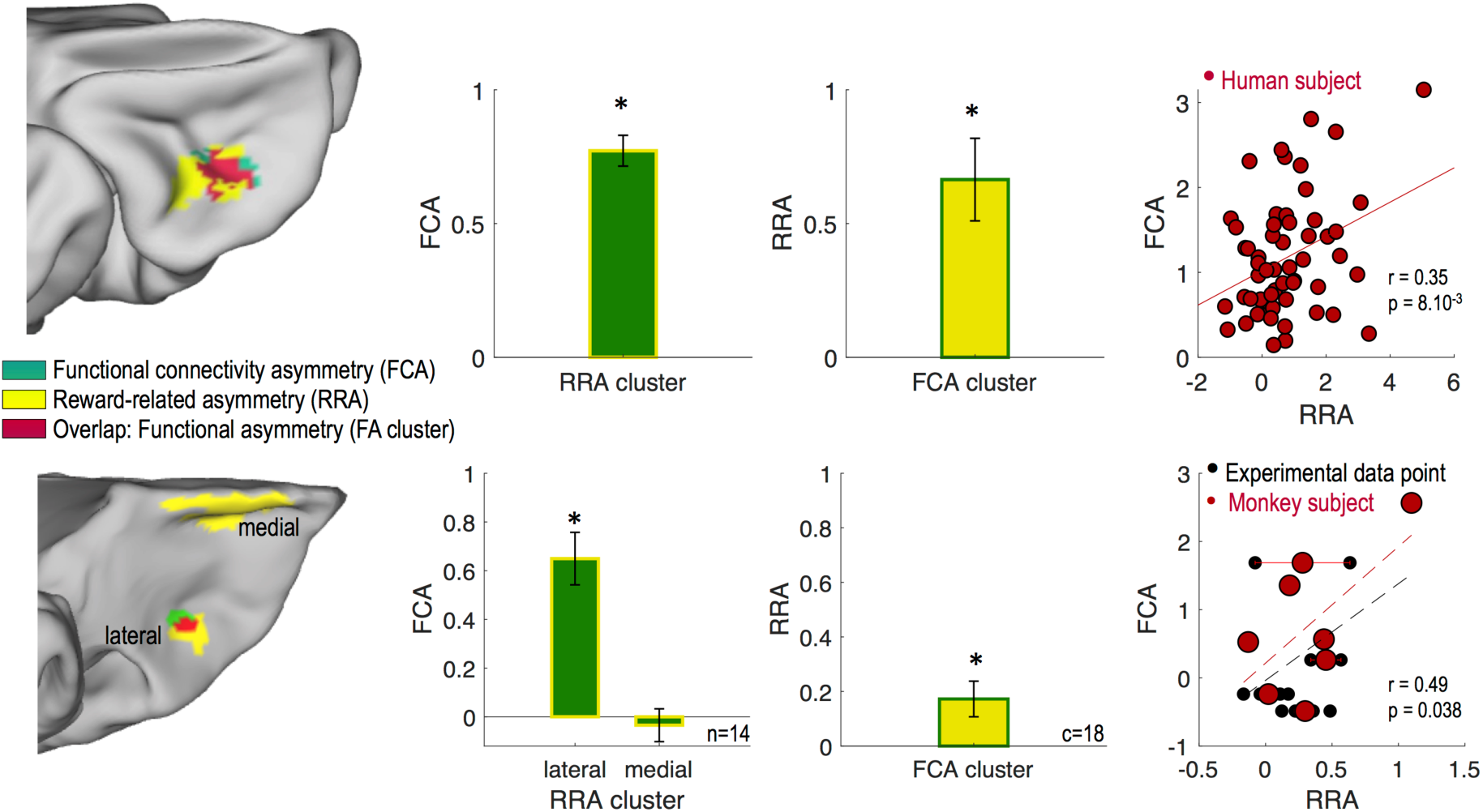
Functional asymmetry in the human and macaque OFC. **First column**. Overlap (red) between the clusters of asymmetry identified in the functional connectivity (FCA, green) and in the reward response (RRA, yellow) analyses on the OFC surface in humans (top) and in macaques (bottom). **Second column**. Mean FCA coefficient across individuals in the RRA clusters (yellow). **Third column**. Mean RRA coefficient across individuals (reward contrasts for monkeys) in the FCA cluster (green). **Last column**. Individual participants’ FCA coefficients plotted as a function of their RRA coefficients in the FA cluster (red). *Top.* Each red point represents one individual. *Bottom.* Each red dot represents one monkey and each black point corresponds to an experimental data point (from 1 to 4 per monkey). Bar plot and error bars represents mean and SEM. Stars indicate significance against 0. n indicate the number of macaques, c indicates the number of contrasts.

Moreover, we extracted the individual participants’ RRA and FCA coefficients from the OFC cluster resulting from the conjunction of the two asymmetry analyses (labeled ‘Functional Asymmetry cluster’ or FA cluster). We found that the two measures of asymmetry, based on RRA and FCA, were strongly correlated in humans (r=0.35, p=8.10^-3^). In macaques, in order to increase the statistical power of the analysis, we decomposed the 14 individual RRA points into experimental data points (18 different contrasts from 4 protocols, see table 1) and again found a significant correlation between RRA and FCA measures (r=0.49, p=0.038). Together, these results suggest that asymmetry in functional connectivity might explain asymmetry of results in task-related activity in both species.

**Table 1.** Monkey ID and protocol details

| monkey ID | rs-MRI | fMRI | protocol ID | Number of sessions | Number of contrasts of interest |
| --- | --- | --- | --- | --- | --- |
| 1 | 1 | 0 | - | - | - |
| 2 | 1 | 0 | - | - | - |
| 3 | 1 | 1 | 4 | 12 | 2 |
| 4 | 1 | 1 | 1 | 10 | 1 |
| 5 | 1 | 1 | 1 | 10 | 1 |
| 6 | 1 | 1 | 1 | 10 | 1 |
| 7 | 1 | 1 | 1 | 10 | 1 |
| 8 | 1 | 0 | - | - | - |
| 9 | 1 | 0 | - | - | - |
| 10 | 1 | 1 | 4 | 13 | 2 |
| 11 | 1 | 0 | - | - | - |
| 12 | 1 | 0 | - | - | - |
| 13 | 1 | 1 | 2 | 12 | 2 |
|  |  |  | 3 | 11 | 2 |
| 14 | 1 | 1 | 2 | 11 | 2 |
|  |  |  | 3 | 10 | 2 |
|  |  |  | 4 | 11 | 2 |
| **Total** | **14** | **8** |  | **200** | **18** |

#### Functional connectivity characteristics

Finally, we compared the functional connectivity of the left and right FA cluster with the whole brain in order to characterize their differences. In humans, we observed that the left FA cluster shows a negative functional connectivity with a network including anterior cingulate cortex (ACC), posterior cingulate cortex (PCC), and temporoparietal junction (TPJ). We will refer to this network as the Default Mode Network (DMN). We also observed that both seeds were positively connected to a frontoparietal network, which we refer to as the Executive Network (ExN, Figure 4). To quantify this difference, we extracted the functional connectivity of each seed vertex from the FA cluster with each vertex in the DMN and the ExN, defined from elsewhere (see Methods). Then, we assessed the effect of FA seed hemisphere (left or right), network (DMN or ExN), and network lateralization (left or right) using a 3-factor ANOVA. We found strong main effects of seed, network, and connectivity lateralization (Seed effect: F(1,228)=53.22, p=1.10^-9^, Network effect: F(1,228)=87.40, p=5.10^-13^, connectivity lateralization: F(1,228)=21.61, p=2.10^-5^), all three 2-factors interaction were also significant (Seed x Network: F(1,228)=20.72, p=3.10^-5^, Seed x connectivity lateralization: F(1,228)=27.73, p=2.10^-6^, Network x connectivity lateralization: F(1,228)=6.37, p=0.015). The triple interaction was not significant (F(1,228)=2.10, p=0.15). Post-hoc multiple comparison tests revealed that both seeds were more connected to the ExN than the DMN [main effect of network, also confirmed by the post-hoc (Tukey HSD) tests of the Network x Connectivity lateralization interaction], but that the left seed was less connected to the DMN compared to the right seed, with no difference of connectivity with the ExN (left vs right seed contrast in relation to DMN: p=6.10^-8^; left vs right seed in relation to ExN: p=0.49).

**Figure 4 –.**
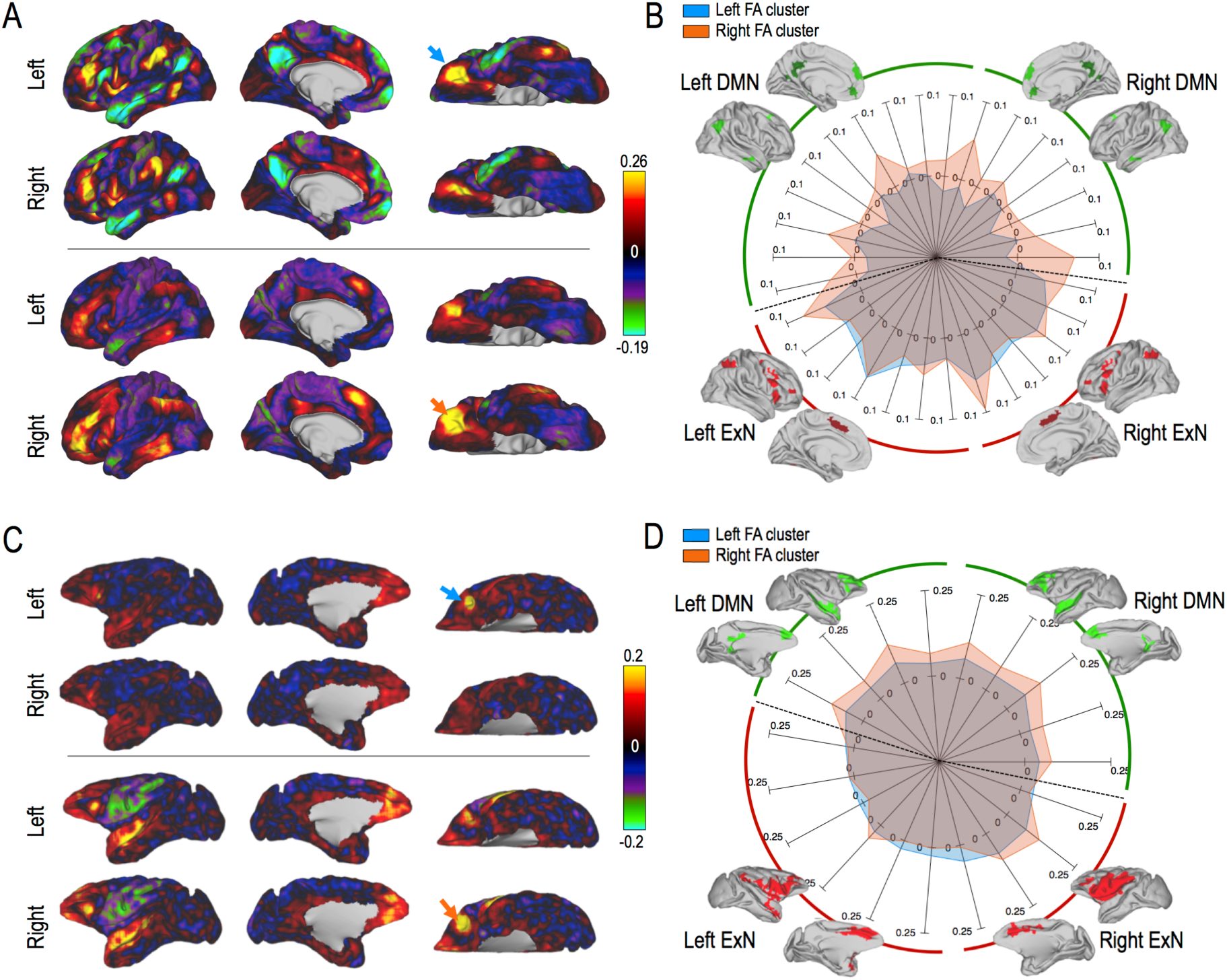
Functional connectivity of the left and right FA clusters. **A.** *Top.* Functional connectivity map of the left FA cluster (blue arrow) with the left hemisphere (first row) and the right hemisphere (second row). *Bottom.* Same as top but for the right FA cluster (orange arrow). Colors indicate the strength of functional connectivity (correlation coefficients). **B.** Connectivity profile (spiderplot) of the left (blue) and right (orange) FA cluster with the Default Mode Network (DMN, green) and the Executive Network (ExN, red). The two networks are decomposed into several subregions that we grouped under the labels ‘Left’ or ‘Right’, i.e. ‘Left DMN’ corresponds to areas belonging the DMN and located in the left hemisphere. Intensities correspond to the coupling of each seed with each target. **C and D** are the same figures as A and B but for macaque data.

In macaques, we observed that the connectivity of the left FA cluster with the rest of the brain was weaker than in the right FA cluster, especially in the DMN. The similar fingerprint analyses revealed results in macaques that were surprisingly similar to those in humans. Indeed, once again, we found main effects of network and connectivity lateralization (Seed effect: F(1,104)=3.63 p=0.07, Network effect: F(1,104)=24.06, p=3.10^-4^, connectivity lateralization: F(1,104)=45.2, p=2.10^-5^), two 2-factors interaction were also significant (Seed x Network: F(1,104)=13.95, p=3.10^-3^, Network x connectivity lateralization: F(1,104)=23.04, p=3.10^-4^, Seed x connectivity lateralization: F(1,104)=1.1, p=0.31). The triple interaction was not significant (F(1,104)=0.69, p=0.42). Post-hoc multiple comparison (Tukey HSD) tests revealed that both seeds were less connected to the ExN than the DMN (main effect of network, also confirmed by the post-hoc tests of the Network x Connectivity lateralization interaction), but the left seed was less connected to the DMN compared to the right seed, with no difference of connectivity with the ExN (left vs right seed in the DMN: p=1.10^-4^; left vs right seed in the ExN: p=0.21). Results are summarized in Figure 4.

#### Morphological characteristics in humans

Given the richness of the HCP data, we were able to further explore some morphological features of the asymmetric OFC FA cluster. We checked whether it was characterized by particular morphological features and found no specific pattern of myelination, gyrification (curvature) or cortical thickness (Figure 5). We compared such features in the left and right FA cluster and found that the myelination of the right FA cluster was higher than in the left FA cluster (t(56)=3.7, p=5.10^-4^). The other features were not significantly different (curvature: t(56)=0.92, p=0.36; cortical thickness: t(56)=-0.94, p=0.35). Although there was an asymmetry in myelination profile, individual variation in the myelination profile asymmetry was not significantly correlated with the RRA, FCA, or FA (mean of RRA and FCA) measures (all p>0.2). The other morphological feature asymmetry coefficients were also uncorrelated with the functional asymmetry measures (all p>0.01, threshold for multiple comparisons). Thus, we found no evidence that the morphological differences in the left and right FA clusters are driving the functional asymmetry observed in that particular area.

**Figure 5 –.**
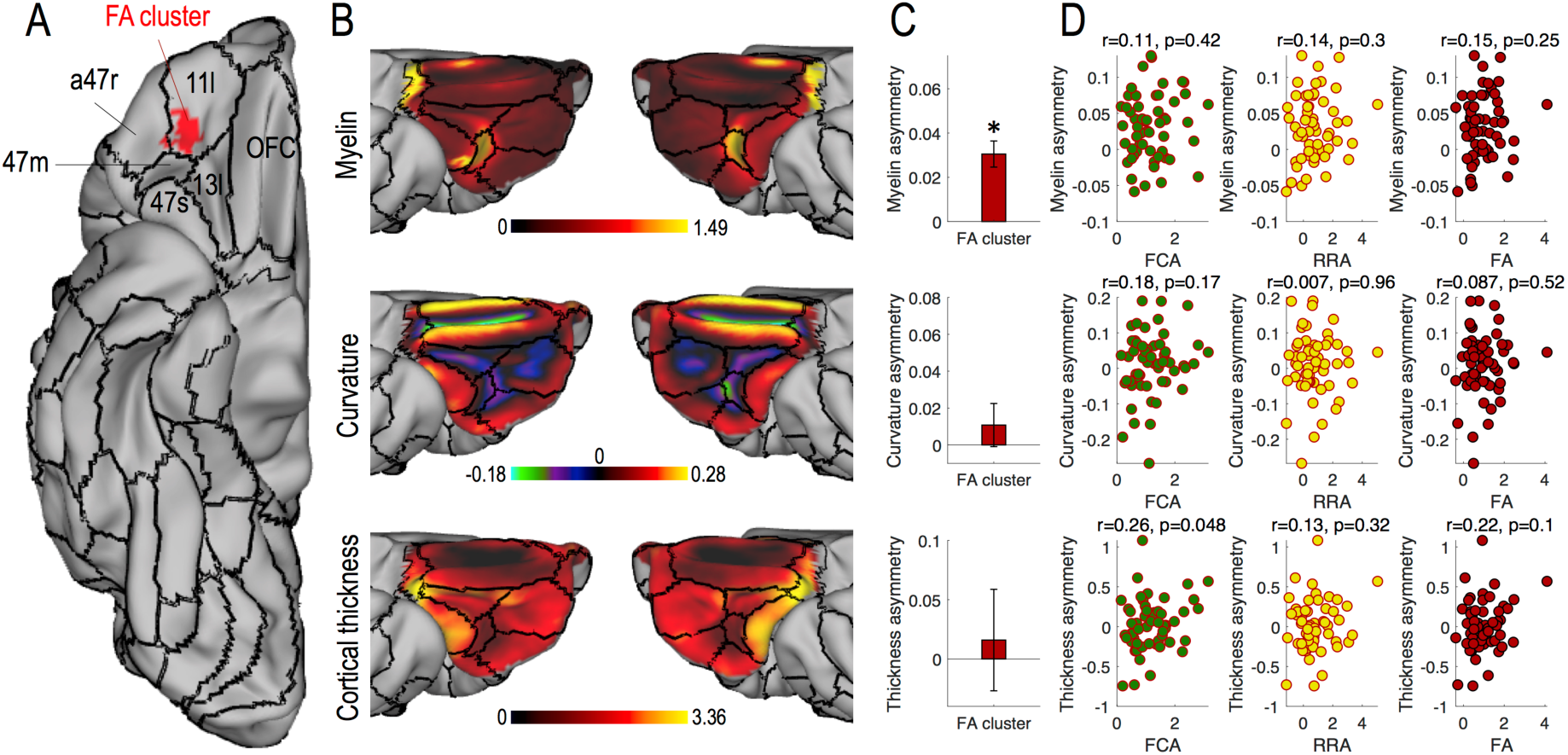
Morphological characteristics of the human FA cluster. **A**. Overlap of the FA cluster (red) and the parcellation from Glasser et al 2016 (29) (black borders). **B**. Morphological features of the OMPFC: Myelin, Curvature (negative in sulci, positive on gyri) and cortical thickness. **C**. Signed difference between left and right morphological features. Star indicates significance against 0. Bar represent the mean across subjects and error bars represent SEM across subjects. **D**. Morphological asymmetries in function of FCA (green), RRA (yellow), and the average of the two measures (FA, red) in the FA cluster.

## Discussion

In the present study, we provide evidence for functional lateralization in OFC. Lateralization in the frontal cortex has been considered most often in relation to language processes and praxis (30–32) but also linked to attention (33) and emotional regulation (34). Although the adaptive consequences of lateralized functions are not well understood, it is thought that hemispheric specialization could increase processing abilities by reducing bilateral redundancy (20). Reward-related asymmetry in the OFC is consistent with many previous studies reporting unilateral responses in the OFC (35–41), there has only rarely been acknowledgement that this is the case (42, 43). Crucially we show an interrelationship across subjects between the reward related asymmetry (RRA) and a functional connectivity-related asymmetry (FCA). Differences between connectivity patterns in the left and right OFC are notably related to their coupling with a set of brain regions often referred to as the DMN. The right OFC was found to be more strongly connected to the DMN than the left OFC. In addition, we observed a similar functional lateralization in the OFC in non-human primates. This result suggests that this asymmetry could have been present in the last common ancestor of humans and old-world monkeys around 29 million years ago. A recent study found an inter-hemispheric OFC asymmetry in rodents in a reversal learning task (44), with the right OFC being more recruited in the task than the left OFC. In tandem with the current results this suggests that reward-related asymmetry in or near OFC might have been a feature of the mammalian brain present since the last common ancestor of rodents and primates more than 100 million years ago.

To our knowledge, this is the first time that the functional asymmetry of the OFC response to reward has been investigated in relation to the same region’s asymmetrical functional connectivity, in both humans and macaques. The reward gambling task used in humans as part of the HCP has some limitations; the simple condition contrast “reward vs punishment” is not ideal for investigating finer aspects of the reward representation. It is therefore difficult to interpret the impact of this OFC lateralization on cognitive processes and behavior. It is possible that the results of studies employing causal approaches such as stimulation or investigation of the effect of brain lesions that have also noted differences in effects in the two hemispheres (13, 14, 45) reflect the same underlying asymmetry as investigated here.

It should, nevertheless be remembered that some studies have reported no effect of OFC lesion laterality (46) or a bilateral OFC responses to reward (47, 48). Therefore, it is important to mention that we do not claim an absolute and total functional dissociation between left and right OFC but rather a graded difference between the contributions that they make. If that is the case, then lateralization in reward-related processing in OFC would resemble lateralization in the language system. It is possible that the relative contribution of each hemisphere’s OFC might differ depending on the requirements of the experimental paradigm. For instance, some studies only report the left OFC to represent outcome information (20), while others only report the right OFC to respond to identity-specific value (19, 20).

It may be worth noting that in our study reward responsivity was investigated in the context of decision making. Intriguingly a recent meta-analysis of lateralization of function suggested that decision-making rather emotion, communication, or perception/action is associated with the OFC lateralization (16). Intracranial electrophysiological recordings in humans have shown that risk-taking biases are driven by a lateralized push-pull neural response, with an increase of high frequency activity in the right hemisphere biasing subjects toward risky bets (43). Alternatively it has been suggested that OFC lateralization might be considered within an exploration/exploitation framework (38). One possibility might be that OFC lateralization is associated with the valence of feedback but no evidence has been found that this is the case (38).

Given that connectivity constrains and partly determines the functions that could be supported by a given brain region (49), one might use rs-fMRI results to further speculate about the nature of the functional differences between the left and right OFC. DMN has been shown to strongly overlap with the social brain network (50). However, responses to social feedbacks, if anything appear stronger in the left OFC than in the right OFC (51). DMN has also been associated with self-referential mental activity, and recollection of prior experiences (52). It might therefore be hypothesized that that it is an internally driven valuation process, i.e. a value assignment that requires individuals to remember or simulate (such as the taste of a cake), that underlies right OFC lateralization. On the other hand, a valuation process linked to external features such as color combination in a painting could recruit the left OFC more. Future investigations will aim at testing this specific hypothesis.

In summary, OFC lateralization has been overlooked or mentioned only in passing in many functional studies. Here, we provide evidence for lateralization in terms of reward-related function and in terms of functional connectivity both in humans and in macaques. Therefore, we strongly encourage future studies to report relative variation in activation in the left and right OFC, and to take into account differences between hemispheres when interpreting the results in OFC.

## Methods

### Subjects

#### Humans

The data used in this study are released as part of the Human Connectome Project (WU-Minn Consortium: Human Connectome Project, RRID: SCR_008749, http://db.humanconnectome.org) (51). We selected the S900 subject release with 7T structural and resting-state MRI (rs-MRI) data. The data were preprocessed according to the HCP pipeline (52). Of the 73 subjects in this specific HCP release, 16 subjects were excluded because of family ties with other subjects in the database. The data analysis was therefore based on 57 subjects (37 females).

Analyses were conducted on the data aligned using areal-feature-based registration (called “MSMAll” for “Multimodal Surface Matching” (29)). This procedure aligns vertices on the cortical surface across subjects not only according to gross folding morphology, but also takes into account the subject-specific functional features, such as the location and distribution of resting-state networks. The MSMAll approach dramatically improves the functional alignment of cortical areas over and above registration based solely on volumetric or surface-based morphological registration. This type of registration is referred to as “area-based” registration and is sometimes considered a near optimal functional alignment (29).

#### Macaques

14 rhesus monkeys (Macaca mulatta, 13 males) were involved in the study. They weighed 7-14 kg and were of 7-13 years of age. They were group housed and kept on a 12hr light dark cycle, with access to water 12-16hr on testing days and with no restriction of access on non-testing days. All procedures were conducted under licences from the United Kingdom (UK) Home Office in accordance with the UK The Animals (Scientific Procedures) Act 1986 and with European Union guidelines (EU Directive 2010/63/EU). Among the 14 monkeys, 8 participated in 4 different experimental tasks (Protocols). The detail of assignment of monkeys to the different tasks is described in table 1.

### Experimental tasks

#### Gambling task in humans

Reward-related BOLD signal was recorded with fMRI during a card-guessing gambling task played for monetary reward that has been previously described (26). Participants completed a card-guessing game where they were required to guess the number (ranging from 1 to 9) on a mystery card in order to win or lose money. Participants were instructed to guess if the unknown card number was more or less than 5 by pressing one of two buttons on a response box. Feedback was given as the revealed card number with a cue to inform the participants if they received a monetary reward, monetary loss or nothing (neutral no reward/loss outcome received for number 5) trial. The task was presented in blocks of eight trials that were either mostly rewarded (six reward trials pseudo-randomly interleaved with neutral and/or loss trials) or mostly loss (six loss trials interleaved with reward and/or loss trials). For each of the two runs, there were two mostly reward and two mostly loss blocks, interleaved with four fixation blocks (15 s duration).

#### Protocol 1 in monkeys: Object Discrimination Reversal Task

The experimental task used in Protocol 1 is described in detail elsewhere (39, 53). Briefly, the task was designed to investigate contingent learning mechanisms and specifically how and where in the brain associations between choice options and outcomes (i.e. reception of reward) resulting from choosing them are formed. Four macaques had to choose between pairs of abstract visual stimuli while in the magnetic resonance imaging (MRI) scanner. On each trial, the two stimuli available for choice (available options) were drawn from a set of three, each associated with distinct reward probabilities. The rewards were delivered probabilistically in a manner that fluctuated across the session, with two of the options reversing toward the middle of a session. Each stimulus’ reward probability was uncorrelated from that of the others. On each trial one of the two available options was chosen by the monkey, the other was unchosen and a third option was invisible and unavailable for choice. In our study, we focused on the receipt of the reward.

#### Protocol 2 and 3 in monkeys: Decision to act Task

The experimental task used in Protocol 2 is described in detail elsewhere (54). Briefly, the task was designed to investigate how contextual factors and internal state, shaped by present and past environment are integrated to influence whether and when to act. 4 monkeys initially performed this task but we only included the two monkeys (13 and 14) who also performed the resting-state fMRI data acquisition. In that task, macaques were trained to track the number of dots on a screen while in the MRI scanner. Dots appeared one at a time on a screen and animals could decide to make a response, at a time of their choice, by tapping on a response pad in front of them. The number of dots on the screen at the time of response determined the probability of reward. Reward probability was drawn from a sigmoid function: the longer the animals waited before responding, more dots appeared on the screen, and the higher was the probability of reward. Different levels of reward magnitude were associated with different dot colors, and the reward magnitude varied from trial-to-trial. Once the monkeys responded, they received drops of juice or no juice according to the reward probability distribution and the time of their response. There was a 4 second delay between the response and the outcome. In the context of our study, two events on each trial were of special interest: the onset of the stimulus (dots), since the color is indicating the expected level of reward, and the outcome (0, 1, 2 or 3 drops of blackcurrant juice).

Data from protocol 3 has not been published yet. However, the task is exactly the same except that the frequency of all the good offers increased and of all the bad offers decreased (i.e., there were more trials with high reward magnitude and less trials with low reward magnitude in protocol 3 compared to protocol 2).

#### Protocol 4: Stimulus-reward association task

The data and results from the experimental task used in Protocol 4 have not been published yet. Briefly, the control task used here investigated how a monkey would respond to visual cues indicative of how much reward could be obtained, or lost (i.e. poured into a visible plastic jar). 4 male rhesus macaques were trained to associate a set of 10 stimuli with various reward magnitudes (i.e. from 0 to 2 drops of reward smoothie that could be either obtained or discarded). On any trial one stimulus was presented on the screen. The monkey had 10 seconds to respond by putting his hand over a homemade infrared sensor. Once selected the stimulus was replaced by a hollow white frame. After a 3.5 to 4.5s delay, the stimulus was presented back (feedback) and the reward delivered. If the monkey did not respond within 10 seconds, the trial was aborted and the same stimulus was presented again after the inter-trial interval. The stimulus-outcome association was probabilistic. In 24% of the trials, the feedback was different from the cue. The obtained reward was always congruent with the displayed feedback.

#### fMRI data acquisition, processing and analysis in humans

The preprocessed 3T data were downloaded from the HCP website for the 57 selected subjects. For each subject, the fMRI data were preprocessed using the HCP functional pipeline, including the volume and MSMAll surface pipeline outputs, motion parameters and FMRIB’s Software Library (FSL, RRID: SCR_002823) (55) files for higher analysis. All preprocessing steps and preliminary analysis are described in (26). Briefly, the HCP ‘fMRIVolume’ pipeline performs gradient unwarping, motion correction, fieldmap unwarping and grand mean intensity normalization on the four-dimensional (4D) time series. These volumes are segmented (Brain Boundary Registration), registered to the T1 anatomical volume using nonlinear transformation (FNIRT) and warped to standard (MNI152) space. Parameter estimates were estimated for a pre-processed time series using a general linear model (GLM) using FMRIB’s improved linear model (FILM) with autocorrelation correction. Predictors were convolved with a double gamma canonical hemodynamic response function to generate regressors. Temporal derivatives of each regressor were added to the GLM as covariates of no interest. Parameter estimates (BOLD) for the contrast (reward > punishment; cope6.feat) were available for 57 participants. We chose this contrast to establish relationships with reward. As the paradigm was a card-guessing task, the contrast corresponded to reward receipt and did not include an anticipation phase.

To obtain group statistics, second level (group) analysis on volumes was conducted using FLAME (FMRIB’s Local Analysis of Mixed Effects) stage 1, part of FSL (version 5.0.8 http://fsl.fmrib.ox.ac.uk/). The main contrast of interest, “Reward versus Punishment”, of each participant was entered into a second level random-effects analysis using a one-sample t-test. The main effect images are all cluster-corrected results with the standard threshold of z>2.3.

For clarity in the data visualization and for a better visual comparison with restingstate data, we then projected the volume result on the averaged MSMAll midthickness surface of all participants, using the ‘wb command’ and ‘volume to surface mapping’ functions from the connectome-workbench (https://www.humanconnectome.org/software/connectome-workbench.html).

To test the asymmetry of reward-related activity, each individual z-stat map corresponding to the ‘reward vs punishment’ contrast was projected onto its corresponding MSMAll surface. Then, the left and right data were extracted from each hemisphere in the OMPFC. The individual unsigned difference between the left and right z-statistics in the OMPFC were computed and then assessed for significance at the group level using permutation tests (see below).

#### fMRI data acquisition and processing in macaques

Awake-animals were head-fixed in a sphinx position in an MRI-compatible chair (Rogue Research, MTL, CA). MRI was collected using a 3T horizontal bore MRI clinical scanner and a four-channel phased array receive coil in conjunction with a radial transmission coil (Windmiller Kolster Scientific Fresno, CA). Each loop of the coil had an 8cm diameter, which ensures a good coverage of the animal’s head. Similar coils have been previously used for awake fMRI studies in primates (39, 56, 57). The chair was positioned on the sliding bed of the scanner. The receiver coils were placed on the side of the animal’s head with the transmitter placed on top. An MRI-compatible screen (MRC, Cambridge) was placed 30cm in front of the animal and the image was projected on the screen by a LX400 projector (Christie Digital Systems). Functional data were acquired using a gradient-echo T2* echo planar imaging (EPI) sequence with a 1.5 x 1.5 x 1.5 mm resolution, repetition time (TR) 2.28 s, echo time (TE) 30 ms and flip angle 90°. At the end of each session, proton-density-weighted images were acquired using a gradient-refocused echo (GRE) sequence with a 1.5 x 1.5 x 1.5 mm resolution, TR 10 ms, TE 2.52 ms, and flip angle 25°. These images were later used for offline MRI reconstruction.

Preprocessing was performed using tools from FMRIB Software Library (FSL) (58), Advanced Normalization Tools (ANTs; http://stnava.github.io/ANTs) (59), Human Connectome Project Workbench (60) (https://www.humanconnectome.org/software/connectome-workbench), and the Magnetic Resonance Comparative Anatomy Toolbox (MrCat; https://github.com/neuroecology/MrCat). First, T2* EPI images acquired during task performance were reconstructed by an offline-SENSE method that achieved higher signal-to-noise and lower ghost levels than conventional online reconstruction (61) (Offline_SENSE GUI, Windmiller Kolster Scientific, Fresno, CA). A low-noise EPI reference image was created for each session, to which all volumes were non-linearly registered on a slice-by-slice basis along the phase-encoding direction to correct for time-varying distortions in the main magnetic field due to body and limb motion. The aligned and distortion-corrected functional images were then non-linearly registered to each animal’s high-resolution structural images. A group specific template was constructed by registering each animal’s structural image to the CARET macaque F99 space (61). Finally, the functional images were temporally filtered (high-pass temporal filtering, 3-dB cutoff of 100s) and spatially smoothed (Gaussian spatial smoothing, full-width half maximum of 3mm).

#### fMRI data analysis in macaques

To perform whole-brain statistical analyses we used a univariate generalized linear model (GLM) framework as implemented in FSL FEAT (62). At the first level, we constructed a GLM to compute the parameter estimates (PEs) for each regressor. The GLMs were constructed based on the specific questions raised in each protocol:

– GLM1 (Protocol 1): *BOLD = βθ + βl DEC + β2 choV + β3 uncV + β4 unpV + β5 choT-uncT + β 6 unpCT + β7 locT + <u>**β8 REW + β9 NOREW**</u> + βl0 c_clo_ + β11 rewTreward +β 12 rewTnoreward + βl 31 e ftuncon v + βl 4 r ightuncon v + ε*
– GLM2 (Protocol 2): *BOLD* = *β0* + *β1* STIM + <u>***β2 expectedReward***</u> + *β3 dotSpeed* + *β4 IT I* + *β*5 *pαstRew* + *β6 pαstαctTime* + *β7 αctTime* + *β8 time* + *β9 rightconv* + *β10 leftconv* + *β11* REW + <u>***β*12 *levelOut***</u> + *β*13 *rightunconv* + *β*14 *leftunconv* + *β*15 *mouth*
– GLM3 (Protocol 3): *BOLD = β*0 + *β*1 STIM + <u>***β*2 *expectedReward***</u> + *β*3 *dotSpeed* + *β*4 *ITI* + *β*5 *pαstRew* + *β*6 *pαstαctTime* + *β*7 *αctTime* + *β*8 *time* + *β*9 *rightconv* + *β*10 *leftconv* + *β*11 REW + <u>***β*12 *levelOut***</u> + *β*13 *rightunconv* + *β*14 *leftunconv* + *β*15 *mouth*
– GLM4 (Protocol 4): *BOLD = βθ + βl DEC + β2 M?ssedDEC + β3 ResponseT?me + β4 decisionHand + <u>**β5 expectedReward**</u>+β6 expectedRewardThrown <u>**+β7 IevelOut**</u> + β8 rewardThrown + β9 RPE + βl 0 RPEThrown + βl 1 leftuncon v + βl 2 ríghtuncon v + βl 3 mouth*

Regressors of interest:

– REW and NOREW: constant regressors were time-locked to onset of feedback, for receipt or non-receipt of the reward
– expectedReward: parametric regressor with up to four levels (depending on protocol), which represents expected reward magnitude
– levelOut: parametric regressor with three or four levels representing the reward outcome on the current trial

Regressors of non-interest:

– STIM: unmodulated regressor representing the main effect of stimulus presentation on responded trials
– DEC: unmodulated decision constant regressor time-locked to onset of the decision
– MissedDEC: unmodulated constant regressor for missed trials in protocol 4
– Cclo: choice location
– choV: chosen option value
– uncV: unchosen option value
– unpV: unpresented option value
– unpCT: unpresented option choice trace
– choT-uncT: choice traces difference between chosen and unchosen options
– rewTreward and rewTnoreward: reward trace when reward is received or not received
– dotSpeed: parametric regressor with 3 levels, representing speed of dots ITI: parametric regressor with 3 levels, representing inter-trial-interval on the current trial
– pastRew: parametric regressor with four levels representing the reward outcome on the past trial.
– pastactTime: actTime on the past trial
– actTime: time-to-act (number of dots at response) on the current trial
– time: parametric regressor representing the time passed since the beginning of the scanning session and locked to the trial onset
– ResponseTime: parametric regressor representing the response time
– decisionHand: parametric regressor representing the hand used to respond
– expectedRewardThrown: parametric regressor with four levels representing the expected amount of reward to be thrown
– rewardThrown: parametric regressor with four levels representing the amount of thrown reward
– RPE: Reward Prediction Error
– RPEThrown: Prediction error on the thrown reward
– Rightunconv and leftunconv: unconvolved categorical regressors for leftwards and rightwards responses
– rightconv and leftconv: convolved categorical regressors for leftwards and rightwards responses
– mouth: distortion due to mouth movements

Regressors in bold are the contrasts linked to reward that we included in our analyses. For each protocol and each contrast, the first-level z-statistics of each session in every monkey were extracted to compute the main effect of reward (fixed effect analysis on volumes). Then, each z-statistic volume was projected onto left and right surfaces and used to compute the asymmetry of reward representation in the OMPFC (linear mixed-effect models that include random factor for protocol and monkeys).

#### rs-MRI data acquisition and processing in humans

The preprocessed 7T data were downloaded from the HCP website. We selected the package called ‘Resting State fMRI 1.6mm/59k FIX-Denoised (compact)’, which contained 1.6mm resolution data. The rs-fMRI acquisitions (including the use of leading-edge, customized MRI hardware and acquisition software) and image processing are covered in detail in (60, 63, 64). After image preprocessing (primarily using the FMRIB Software Library, FSL, RRID:SCR_002823) (58), FreeSurfer (RRID:SCR_001847) (65), and Connectome Workbench (66) software packages), the functional timeseries are filtered and artefacts are removed using an automated data-driven approach that relies on ICA decomposition and hand-trained hierarchical classification (FMRIB’s ICA-based X-noisifier [FIX]) (63). We concatenated the MSMAll data from the 4 available resting-state sessions (demeaned then concatenated) to obtain one time series per participant.

#### rs-MRI data acquisition and processing in macaques

The 14 monkeys were scanned under anesthesia to acquire resting-state data. fMRI and anatomical scans were collected according to previously used protocols (67). Anesthesia was induced using intramuscular injection of ketamine (10 mg/kg) combined with either xylazine (0.125–0.25 mg/kg) or midazolam (0.1 mg/kg) and buprenorphine (0.01 mg/kg). Macaques also received injections of atropine (0.05 mg/kg), meloxicam (0.2 mg/kg), and ranitidine (0.05mg/kg). Anesthesia was maintained with isoflurane. Isoflurane was selected because it has been demonstrated that resting-state networks are still present using this agent for anesthesia (68). The anesthetized animals were placed in an MRI-compatible stereotactic frame (Crist Instrument) in a sphinx position within a horizontal 3T MRI scanner with a full-size bore. The same coils as for awake scans (see fMRI data acquisition) were used for data acquisition. Whole-brain BOLD fMRI data were collected using the following parameters: 36 axial slices, resolution of 1.5 × 1.5 mm, slice thickness of 1.5 mm, TR of 2280 ms, TE of 30 ms, 1600 volumes. Structural scans were acquired in the same session using a T1-weighted MP-rage sequence (no slice gap, 0.5 × 0.5 × 0.5 mm, TR of 2500 ms, TE of 4.01 ms and 128 slices).

The detailed preprocessing pipeline for the resting-state fMRI has been described elsewhere (69, 70). Briefly, after reorientation to the same convention for all functional EPI datasets, the first volumes were discarded to ensure a steady radio frequency excitation state. EPI timeseries were motion corrected using MCFLIRT (71). Brain extraction, bias-correction, and registration were achieved for the functional EPI datasets in an iterative manner, the mean of each functional dataset was registered to its corresponding T1w image using rigid-body boundary-based registration (FLIRT, (71, 72))). EPI signal noise was reduced both in the frequency and temporal domain. The functional timeseries were high-pass filtered with a frequency cut-off at 2000 s. Temporally cyclical noise, for example originating from the respiration apparatus, was removed using band-stop filters set dynamically to noise peaks in the frequency domain of the first three principal components of the timeseries. To account for remaining global signal confounds we considered the signal timeseries in white matter and meningeal compartments, there confound parameters were regressed out of the BOLD signal for each voxel. Following this confound cleaning step, the timeseries were low-pass filtered with a cut-off at 10 s. The data were transformed to F99 and spatially smoothed using a 2 mm FWHM Gaussian kernel. Lastly, the data timeseries were demeaned to prepare for functional connectivity analyses.

#### rs-MRI data analysis

All analyses and statistics were conducted in Matlab 2018b (MATLAB and Statistics Toolbox Release 2017a, The MathWorks, Inc., Natick, Massachusetts, United States, RRID: SCR_001622, www.mathworks.com) with in-house bespoke scripts calling Workbench executables.

Group analyses using MIGP (MELODIC’s Incremental Group-PCA) were first conducted to investigate the global patterns of asymmetry in the orbito-medial prefrontal cortex (OMPFC). MIGP analysis corresponds to a group Principal Component Analysis, as described in (73). The brain activity time series of each participant are sequentially included in a PCA analysis in order to provide a close approximation to the full concatenation of all participant time series, without large memory requirements. The output of this analysis is a time series of similar size to an individual time series.

#### Network definition

In humans, to assess the connectivity of regions of interest to the DMN and to the ExN, the names of the two networks were entered as a topic term in www.neurosynth.org and the association (for the DMN) and uniformity test (for the ExN) maps were downloaded. Maps were then projected onto surfaces and thresholded for clusters bigger than 100 vertices.

In macaques, the networks were defined from the connectivity of bilateral seeds in the anterior cingulate sulcus (DMN) and the mid-cingulate sulcus (ExN).

#### ROI definition

Regions of interest (ROI) were drawn manually on the ventro-medial prefrontal cortex (vmPFC) and the orbitofrontal cortex (OFC), to cover a large portion of the orbito-medial prefrontal cortex (OMPFC). The dorsal medial boundary was delineated by an arbitrary horizontal line that runs from the front of the brain to the genu of the corpus callosum. The ventral surface of the frontal lobe was included from the frontal pole rostrally to the anterior perforated substance caudally.

#### Functional Connectivity Asymmetry coefficient

To investigate the asymmetry of connectivity between the left and the right OMPFC, four measures of asymmetry were used:

- Ipsilateral Functional Connectivity Asymmetry (FCA_Ipsi_): Difference between the connectivity of the left OMPFC (OL) with the left hemisphere (HL) and the right OMPFC (OR) with the right hemisphere (HR).

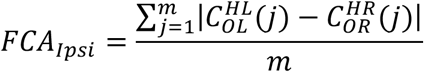
- Contralateral Functional Connectivity Asymmetry (FCA_contra_): Difference between the connectivity of the left OMPFC (OL) with the right hemisphere (HR) and the right OMPFC (OR) with the left hemisphere (HL).

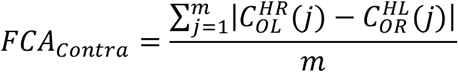
- Left Functional Connectivity Asymmetry (FCA_Left_): Difference between the connectivity of the left OMPFC (OL) with the left hemisphere (HL) and the right OMPFC (OR) with the left hemisphere (HL).

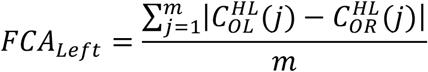
- Right Functional Connectivity Asymmetry (FCA_Right_): Difference between the connectivity of the left OMPFC (OL) with the right hemisphere (HR) and the right OMPFC (OR) with the right hemisphere (HR).

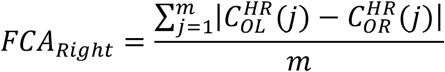

With m the number of vertices on each hemisphere, 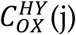 the connectivity of every n vertices of the X (left or right) OMPFC with a vertex j of the ϒ (left or right) hemisphere. FCA is a vector of n elements, visually represented on the heat maps on Figure 2.

#### Statistical assessment

The statistical validity of our results was assessed by extracting variables of interest from each subject and testing for significance at the group level using one-sample t-tests. When assessing significance of clusters on resting-state MRI data, each FCA map was computed for every subject. The main effect was then tested using one-sample student t-test (two-tailed).

To assess the statistical validity of the RRA clusters in both humans and monkeys, we used the Fisher randomization test (74) with 10000 randomizations of the RRA values (z-scored) of each subject. The maximal cluster-level statistics (the sum of t-values across contiguous points passing a significance threshold of 0.01 (z=2.3)) were extracted for each shuffle to compute a ‘null’ distribution of effect size across the OMPFC mask. For each significant cluster in the original (non-shuffled) data, we computed the proportion of clusters with higher statistics in the null distribution, which is reported as the ‘cluster corrected’ p-value (p_corr_) (75).

#### Anatomical MRI data acquisition and analyses

The preprocessed anatomical 7T data were downloaded from the HCP website. We selected the package called ‘Structural Preprocessed for 7T (1.6mm/59k mesh)’, which contained 1.6mm resolution data, collected at 3T. In this package, myelin, curvature and cortical thickness maps are available for each subject, registered on MSM-All, making those maps comparable with the connectivity maps. When investigating the morphological features of the OMPFC, we extracted the values of those maps for each subject and computed the mean of each feature.

## Acknowledgments

Human data were provided by the Human Connectome Project, WU-Minn Consortium (Principal Investigators: David Van Essen and Kamil Ugurbil; 1U54MH091657) funded by the 16 NIH Institutes and Centers that support the NIH Blueprint for Neuroscience Research; and by the McDonnell Center for Systems Neuroscience at Washington University. Funding for this work was provided by the Wellcome Trust (grant nos. 203139/Z/16/Z, WT100973AIA, 103184/Z/13/Z and 105238/Z/14/Z), the Medical Research Council (grant nos. MR/P024955/1 and G0902373), the Bettencourt Schueller Foundation and Christ Church, University of Oxford. We are very grateful for the care afforded to the animals by the veterinary and technical staff at the University of Oxford.

## Author contributions

A.L.-P. and J.S. designed the study. L.R. collected and pre-processed the monkey resting-state data. M.F.S.R. designed the monkey experiments from protocols 1 to 3. J. S., K.M and E.F.F designed the monkey experiment from protocol 4. N.K., D.F, K. M, E.F.F. collected, pre-processed and provided the first-level analyses of the monkey fMRI data. A.L-P. performed all the data analyses in humans and the asymmetry-related analyses in monkeys. A.L.-P., J.S and M.F.S.R. wrote the manuscript. All authors discussed the results and commented the manuscript.

